# Infection with a small intestinal helminth Heligmosomoides polygyrus bakeri consistently alters microbial communities throughout the small and large intestine

**DOI:** 10.1101/575787

**Authors:** Alexis Rapin, Audrey Chuat, Luc Lebon, Mario M. Zaiss, Benjamin Marsland, Nicola L. Harris

## Abstract

Increasing evidence suggests that intestinal helminth infection can alter intestinal microbial communities with important impacts on the mammalian host. However, all of the studies to date utilize different techniques to study the microbiome and access different sites of the intestine with little consistency noted between studies. In the present study, we set out to perform a comprehensive analysis of the impact of intestinal helminth infection on the mammalian intestinal bacterial microbiome. For this purpose, we investigated the impact of experimental infection using the natural murine small intestinal helminth, *Heligmosomoides polygyrus bakeri* (Hpb) and examined possible alterations in both the mucous and luminal bacterial communities along the entire small and large intestine. We also explored the impact of common experimental variables, including the parasite batch and pre-infection microbiome, on the outcome of helminth-bacterial interactions. This work provides evidence that helminth infection reproducibly alters intestinal microbial communities – with an impact of infection noted along the entire length of the intestine. Although the exact nature of helminth-induced alterations to the intestinal microbiome differed depending on the parasite batch and microbiome community structure present prior to infection, changes extended well beyond the introduction of new bacterial species by the infecting larvae. Moreover, striking similarities between different experiments were noted, including the consistent outgrowth of a bacterium belonging to the Peptostreptococcaceae family throughout the intestine.

**Author Summary:** Increasing evidence indicates a role for interactions between intestinal helminths and the microbiome in regulating mammalian health, and a greater understanding of helminth-microbiota interactions may open the path for the development of novel immunomodulatory therapies. However, such studies are hampered by the inconsistent nature of the data reported so far. Such inconsistancies likely result from variations in the experimental and technological methodologies employed to investigate helminth-microbiota interactions and well has natural variation in the starting microbiome composition and/or worm genetics. We conducted a thorough study in which the reproducibility of helminth-induced alterations of microbial communities was determined and impact of common experimental variables – such as the starting microbiome and parasite batch - was determined. Our work reveals the robust ability of small intestinal helminth infection to alter microbial communities along the entire length of the intestine and additionally identifies a single bacterium that is strongly associated with infection across multiple experiments.

## Introduction

The eradication of helminths from developed regions constitutes a major achievement towards the improvement of human health and continued efforts to lower the burden of infection in endemic areas remain essential. Yet intestinal helminths have inhabited the intestine of mammals throughout evolution and are highly likely to have interacted with, and impacted on, the complex bacterial communities present at this site (1). The eradication of helminths is also thought to contribute to the increased incidence of immune disorders, including allergy and autoimmunity, observed in developed societies (2, 3). The mammalian host and its common intestinal inhabitants – including helminths and bacteria - form a complex ecosystem, and the disruption of one of these partners may have important implications for the other players of the ecosystem and ultimately for human health. Indeed evidence is arising indicating that interactions between helminths and bacteria do occur and that these interactions can impact on the mammalian host to shape the host immune response (4, 5). Although the hypothesis that intestinal helminth infection impacts on the mammalian intestinal microbiome is generally accepted, the impact of intestinal helminth infection on the human intestinal microbiome has only been investigated in a small number of studies, and studies using veterinary animals or experimental models have yielded contrasting results. These differences are to be expected as the field has been hampered by difficulties characterizing the intestinal microbiome as a result of strong inter-individual variations not only in humans but also in inbred mice (6). Past studies have also employed a wide array of helminth parasites, host species, and sampling sites and utilized diverse technologies to characterize the microbiome (1), making cross study comparisons difficult.

Many of the experimental studies have utilized the natural murine parasite, *Heligmosomoides polygyrus bakeri* (Hpb), which has a strictly enteric life cycle that is similar to the one of trichostrongyloid parasites affecting human and livestock (7–9). Experimental infection with this parasite occurs via oral gavage of infective larvae from laboratory cultures. Following gavage the larvae penetrate the intestinal epithelium in the upper part of the small intestine and reside in the intestinal outer muscle layer of the intestinal wall (10). Here the worm matures into its sexually mature form in a process that includes two successive moults at days 3-4 and 8-9 post-infection. It finally exits the host tissue and establishes itself as an adult worm in the lumen of the small intestine, where it can survive for months (11, 12). Hpb infection triggers a highly polarized type two immune response characterized by the production of IL-4, IL-5 and IL-13 cytokines, proliferation of IgE and IgG1 producing B cells and mucus secretion (13, 14). Increased smooth muscle contractility together with increased epithelial cells secretions favor the eventual expulsion of adult worms from the intestinal lumen in a combined mechanism that is commonly referred to as the “weep and sweep” response (15)

To date seven studies have investigated the possible interaction between Hpb and the intestinal microbiome (4,5,16–20). The first of these used a method based on the generation of 16S clone libraries and Sanger sequencing to investigate the impact of infection of the bacterial communities and reported higher proportions of Lactobacillaceae in the ileum of infected mice (16). A later study, using real-time PCR and culture-based methods, reported differences between infected and non-infected mice in the bacterial communities residing in the ileum, the cecum and the colon at two weeks post infection (17). Another study using qPCR and Sanger sequencing reported the association of Hpb infection with the abundance of the bacterium *Lactobacillus taiwanensis* in the duodenum of susceptible C57BL/6 mice but not resistant Balb/c mice. These authros also reported that administration of *L. taiwanensis* to resistant Balb/c mice resulted in higher worm burdens, suggesting that L. taiwanensis promotes helminth infection (18). More recently, using high throughput sequencing of the bacterial 16S gene, our own group reported that Hpb infection leads to long-lasting impacts on cecal bacterial communities which are maintained following fecal transfer (4). These findings were confirmed by a later study using a similar method and fecal transfer but focusing on the colonic bacterial communities (19). Lastly, Hpb infection has been shown to provide resistance against *Bacteroides vulgatus* outgrowth in Nod2 deficient mice through a mechanisms involving host type two cytokine production(5). Of note, Hpb excretory/secretory products were recently described to exert anti-microbial activity, raising the possibility that direct helminth-bacterial interactions may take place (20). Taken together these observations endorse the view that intestinal helminths can and do impact on the microbiome, but indicate that the outcome of helminth-microbial-host interactions may be variable.

Numerous factors might be expected to influence the impact of helminth infection on the intestinal microbiome including the genetically variability of the parasites used for infection, the pre-infection or ‘starting’ microbiome of the host, and the possible introduction of bacteria from the infective parasitic larvae. Moreover, Hpb-associated changes in the bacterial community composition may occur only within distinct sites of the intestine, and the infection may impact communities in the intestinal mucus layer differently from the communities present in the intestinal lumen. We considered it particularly important address the latter question given the hypothesis that the worm may impact on the intestinal microbiome indirectly through the host immune response (5), and would thus be expected to have a major impact on those bacteria living in close association with the host intestinal epithelium. We therefore set out to determine the full extent of the impact of Hpb infection on bacterial communities by analyzing microbiomes from the intestinal lumen and epithelium-attached mucus at multiple points along the entire length of the small and large intestine. We also investigated the impact of common variables, including the use of different parasite batches or mice with distinct pre-infection microbiomes, on the ability of Hpb to alter murine intestinal microbial communities.

## Results

### Small intestinal helminth infection impacts on the microbiome present at multiple sites along the small and large intestine, with distinct parasite batches contributing to variability in the exact bacterial community composition found at each site

Many parasitic helminths, including Hpb exhibit a sexual reproductive cycle resulting in large genetic heterogeneity amongst individual worms. Even parasites maintained in laboratories for use in experimental work are produced in batches that can each be considered as different communities and may show significant differences including in term of virulence (11, 21). To test the impact of distinct parasite batches on helminth-induced microbial changes, we assessed the impact of two distinct batches of worms on the same microbiome by performing a large experiment in which all mice were subjected to ‘normalization’ of the ‘starting intestinal microbiome’ by mixing beddings between cages and randomizing mice once a week for four weeks prior to infection (Fig 1). Normalization of the microbiome is necessary as mice bred in the same facility demonstrate inter-cage variations (6). Mice were then sacrificed at day 40 post infection and the bacterial communities in the duodenum, jejunum, ileum, cecum, colon, and feces evaluated by high-throughput sequencing of the v1-v2 hyper-variable regions of the bacterial 16S rRNA gene as described in the Materials and Methods section. Statistical evidences that infection with Hpb affected the composition of the bacterial communities were found at multiple intestinal sites, both in the mucus and in the luminal content (Fig 2). These evidences were reproducibly observed using both batches of Hpb, either in the mucus or the lumen of the duodenum, the jejunum, the cecum and the colon, but not in the ileum (Fig 2). Interestingly, no statistical evidences were observed in the feces with neither of the two batches of worms (Fig 2). In addition, infection with batch a larvae led to a higher diversity in the cecum lumen while infection with batch b larvae led to a higher diversity in the colonic lumen, as shown in term of species richness (Fig 3A) and Shannon diversity index (Fig 3B).

**Fig 1.**
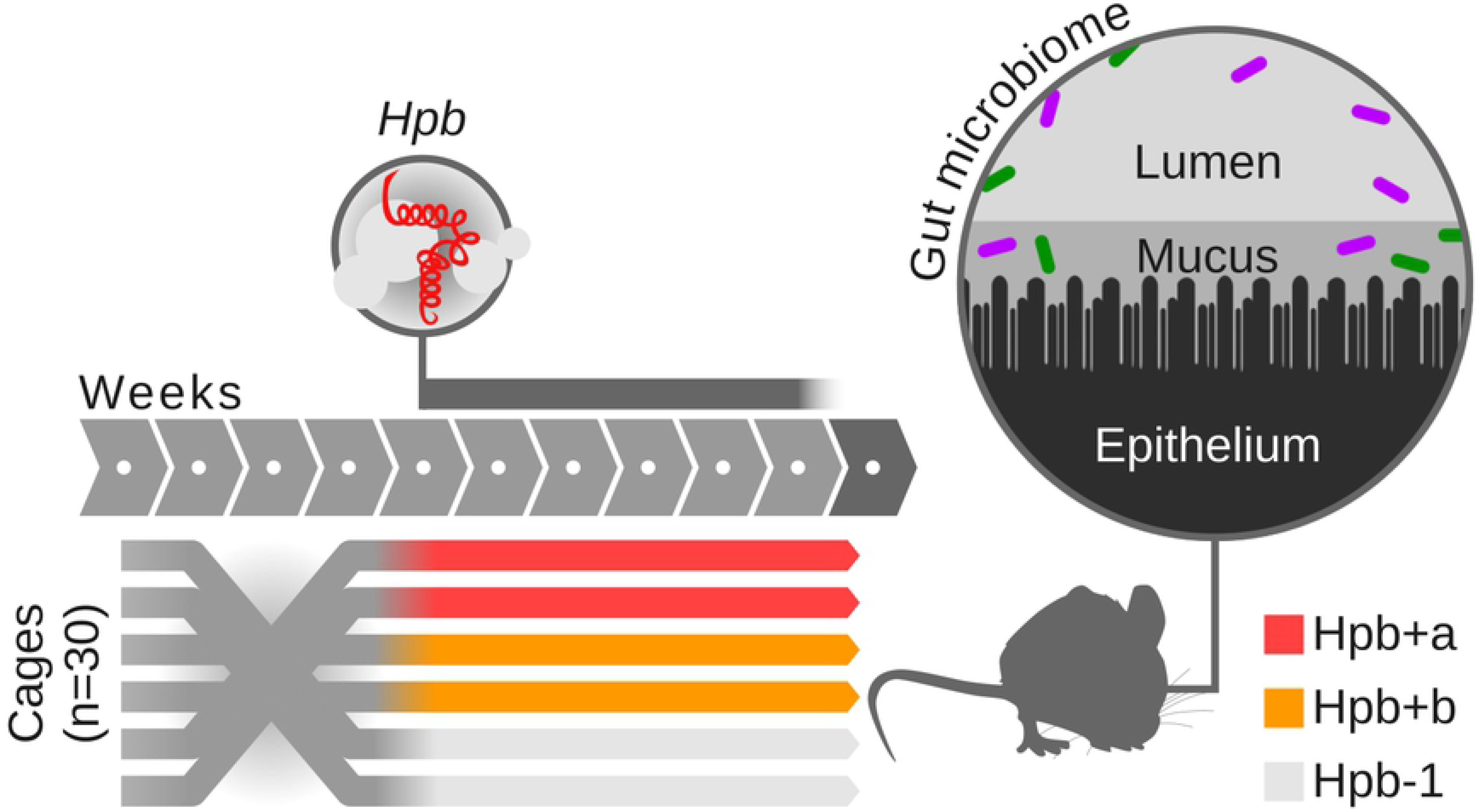
Experimental design of experiment 1 (exp. 1) Analysis of the bacterial communities in the intestinal lumen and intestinal mucus layer 40 days post infection with two distinct batches of Hpb larvae.

**Fig 2.**
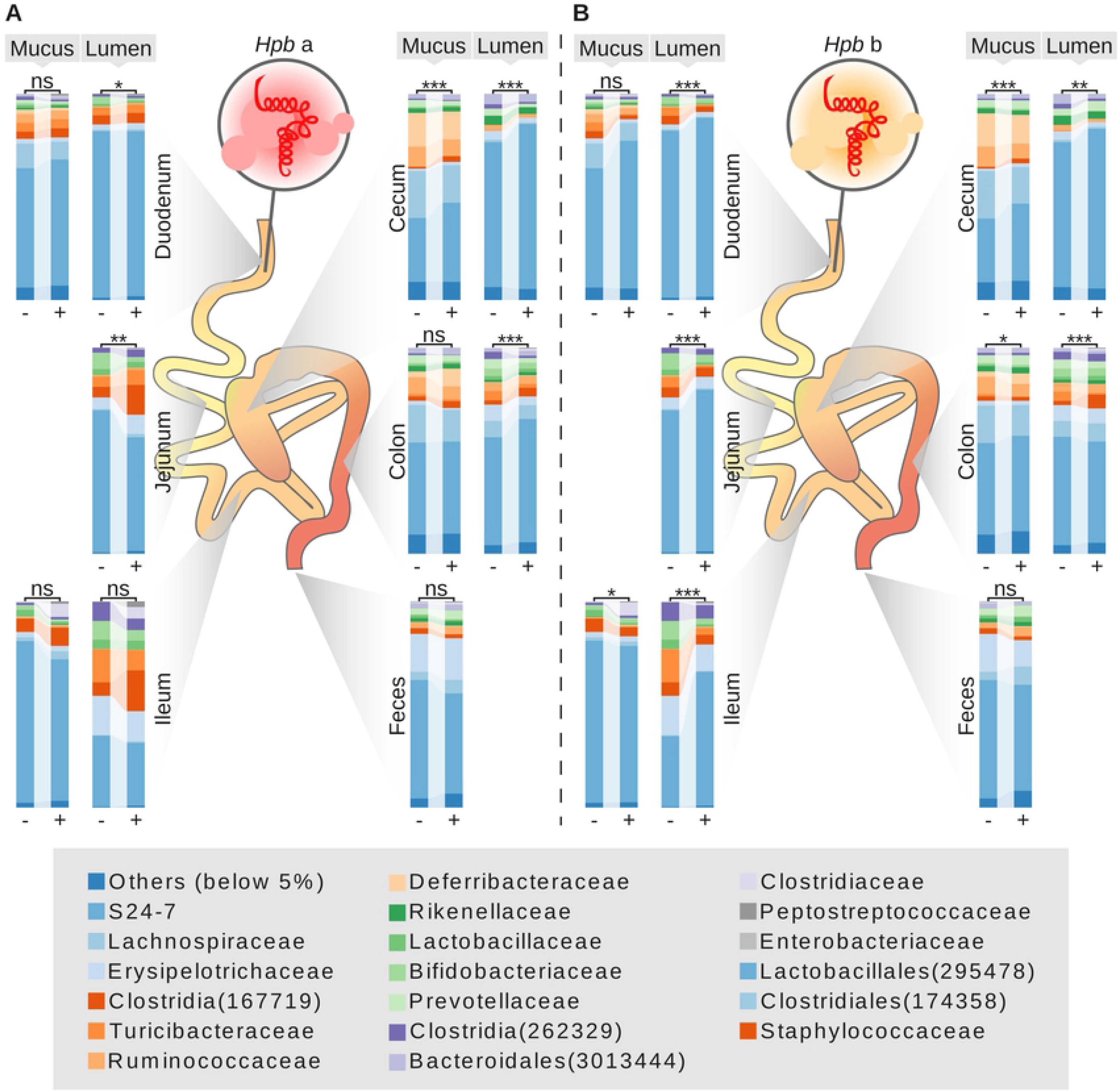
Bacterial communites are affected by Hbp infection at multiple sites along the gut. Average proportions of bacterial families in non-infected (−) and infected (+) mice from exp. 1. (A) Non-infected mice (Hpb-1) are compared to infected mice with infective larvae from batch a (Hpb+a). (B) Non-infected mice (Hpb-1) are compared to infected mice with infective larvae from batch b (Hpb+b). Groups were compared using the Adonis method based on the Jaccard distance (p<0.05: *, p<0.01: **, p<0.001: ***, p>0.05: ns).

**Fig 3.**
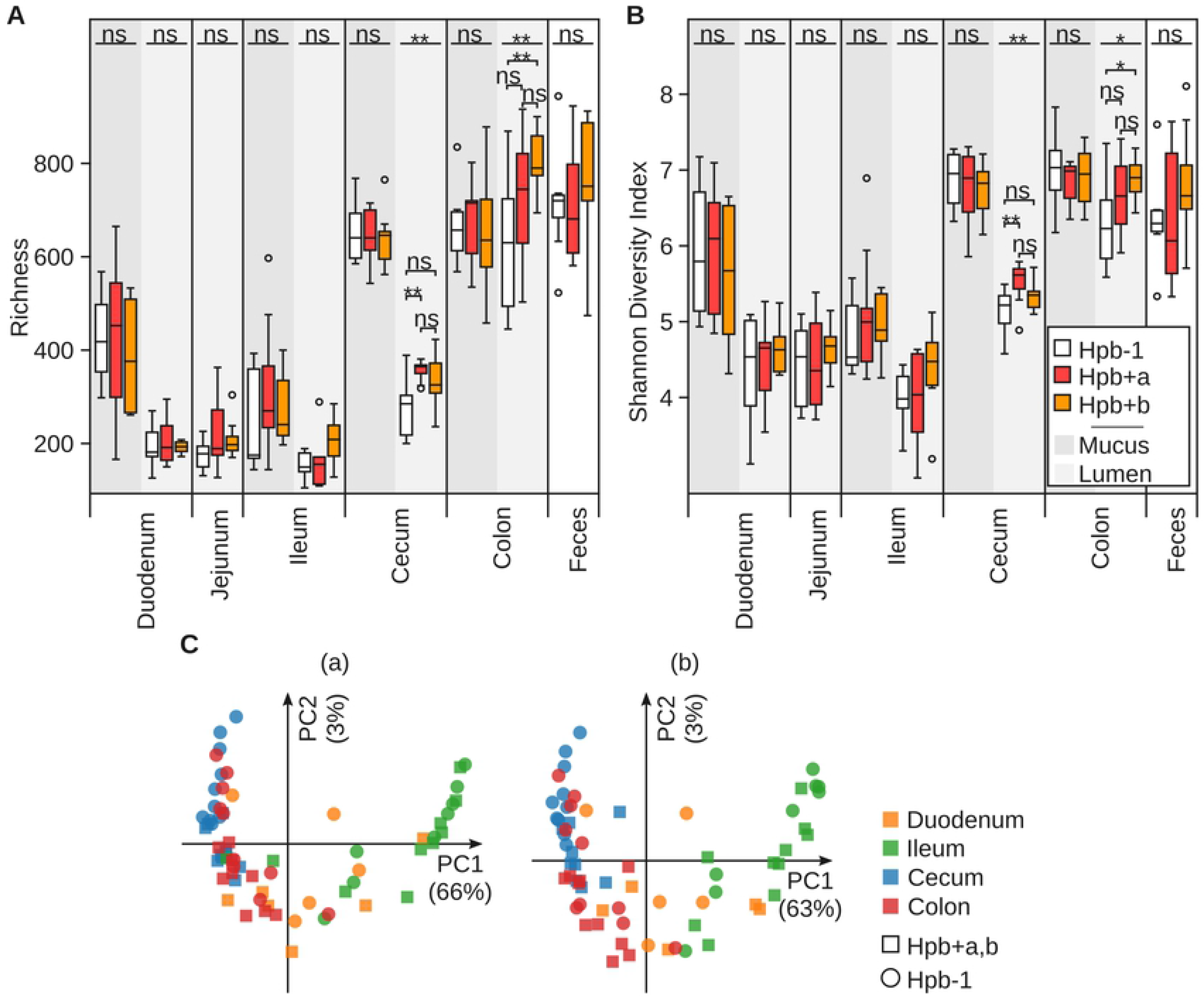
Hpb infection affects bacterial community diversity. Bacterial communities diversity in the mucus layer and lumen of the intestinal tract and feces in terms of richness (A) and Shannon diversity index (B) for non-infected mice from exp. 1 (Hpb-1), infected mice with infective larvae from batch a (Hpb+a) and infected mice with infective larvae from batch b (Hpb+b). Groups were compared by ANOVA (p<0.05: *, p<0.01: **, p<0.001: ***, p>0.05: ns). (C) Principal coordinates analysis (PCoA) based on the Jaccard distance of bacterial communities in the mucus layer along the intestinal tract for non-infected mice from exp. 1 and infected mice with infective larvae from batch a (a) or batch b (b).

Beyond these differences, a PCoA analysis highlighted that bacterial communities tend to cluster according the sampling site rather than infection, with the ileum clearly separating from the rest of the samples (Fig 3C). This indicates that although helminth infection can alter bacterial communities, the niche provided by distinct intestinal sites plays a more important role in determining community structure, with the ileum forming a particularly unique environment as compared to the rest of the intestine. Differences in the proportions of individual bacterial species between Hpb infected and non-infected mice at each sampling site along the intestine notably involved members of the Lachnospiraceae, Clostridiaceae, S24-7, Lactobacillaceae, Ruminococcaceae and Peptostreptococcaceae families, a member of the class Clostridia as well as members of the Allobaculum, Bifidobacterium, Ruminococcus, Sutterella and Turicibacter genera (Fig 4). Finally, the relative abundance of some bacteria were found to be significantly (p<0.05) altered across all studies in response to infection (Fig 5). Most strikingly, a member of the Peptostreptococcaceae family was found in higher proportions in the intestinal mucus layer of infected mice at all sampling sites along the intestine independently of the larvae batch used to infect the mice (Fig 5A and Fig 5C). By contrast, a member of the Bifidobacterium genus was found in lower proportions in both intestinal mucus and lumen at all sampling sites in mice infected with larvae from batch a, but was not detected in mice infected with batch b larvae (Fig 5A and Fig 5B). Membesr of the Allobaculum and Turicibacter genus’s were found in higher proportions in the intestinal lumen of all infected mice, regardless of larval batch used (Fig 5B and Fig 5D).

**Fig 4.**
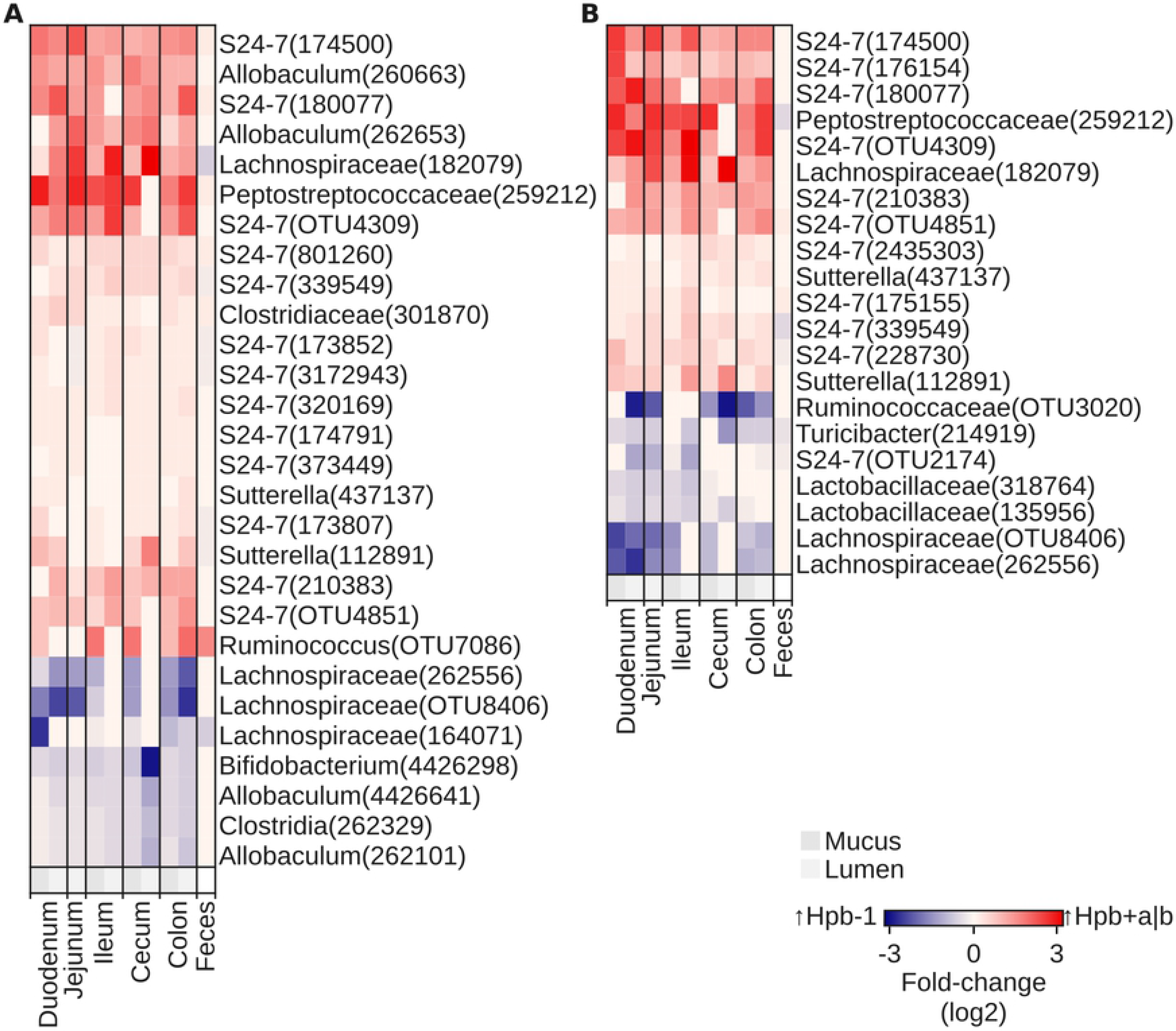
Select bacteria are altered by Hpb infection. Differences in proportions of selected bacterial species in the mucus layer and lumen of the intestine and feces between non-infected mice from exp. 1 (Hpb-1) and mice infected with larvae from batch a (Hpb+a) (A) or batch b (Hpb+b) (B). Differences are represented by the log value of the fold-change as described in the Materials and Methods chapter above. Species were selected based on statistical significance and consistency across multiple sampling sites as described in the Methods section above. Higher proportions in samples from Hpb-1 are represented in a negative (blue) scale while higher proportions in infected samples are represented in a positive (red) scale.

**Fig 5.**
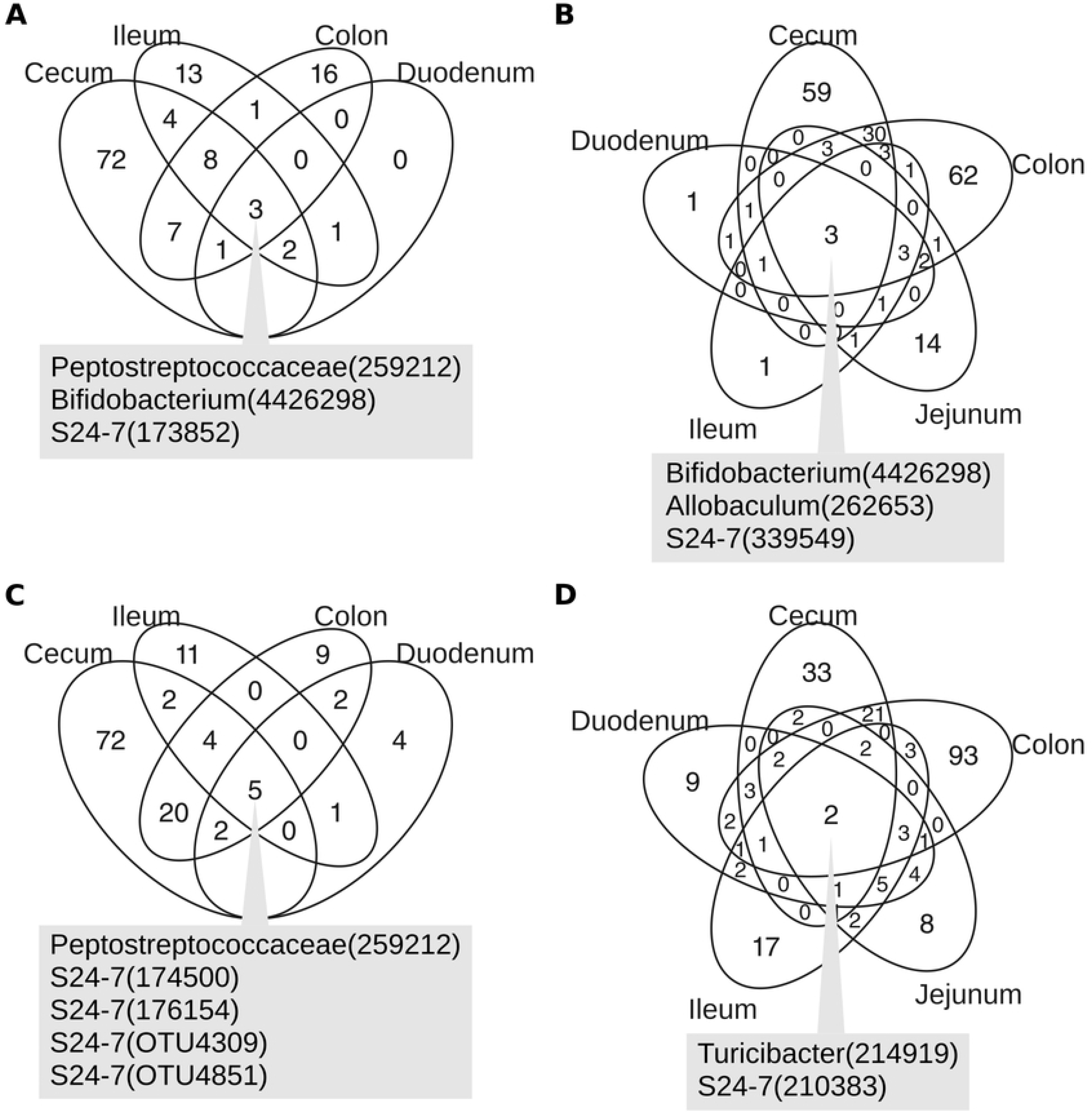
A subset of bacteria are consistently associated with Hpb infection across all intestinal sites. Venn diagrams highlighting overlaps between sampling sites along the intestine for all significant (p<0.05) differences in proportions of observed species between non-infected mice from exp. 1 (Hpb-1) and mice infected with larvae from batch a (Hpb+a) (A, B) or batch b (Hpb+b) (C, D) in the mucus layer (A, C) and the lumen (B, D).

Taken together these data demonstrate that infection with a small intestinal helminth, Hpb, leads to significant changes in bacterial community composition throughout the small and large intestine, but that parasitic variability (resulting from batch variability) can influence the exact nature of these changes.

### The ‘starting point’ of the host intestinal microbiome impacts on the outcome of helminth-bacterial interactions

Beyond inter-individual or inter-cage variations within a given experiment, the microbial communities of mice bred in laboratories are well known to differ between facilities and to change over time (22, 23). For evident reasons, any impact of helminth infection on the microbiome would first depend on its initial state. For example, outgrowth of particular species following helminth infection could only be observed if these species were present in the community prior to the infection. To gain further insight into the robustness of the observed impact of Hbp infection on intestinal bacterial communities, we conducted a second experiment (exp. 2) employing a distinct “starting point” to our first experiment (exp. 1, reported in Figures 1–5). These different “starting points” were obtained by simply conducting experiments at different time periods (approximately 12 months) using mice obtained from the same provider. As for exp. 1 normalization of the starting microbiome within mice entered into exp. 2 was achieved by employing a period where mice were randomized and bedding were mixed between cages once a week for four weeks prior to infection (Fig 6). By necessity the experiment also employed the use of a new batch of parasite larvae (batch c) as Hpb larvae only maintain viability for a period of 8-12 weeks. As expected major differences were found between bacterial communities from the mice employed for exp. 1 (Hpb-1) and exp. 2 Hpb-2 (Fig 7A). Notably, 510 identified OTUs were not shared between these two groups (Fig 7B).

**Fig 6.**
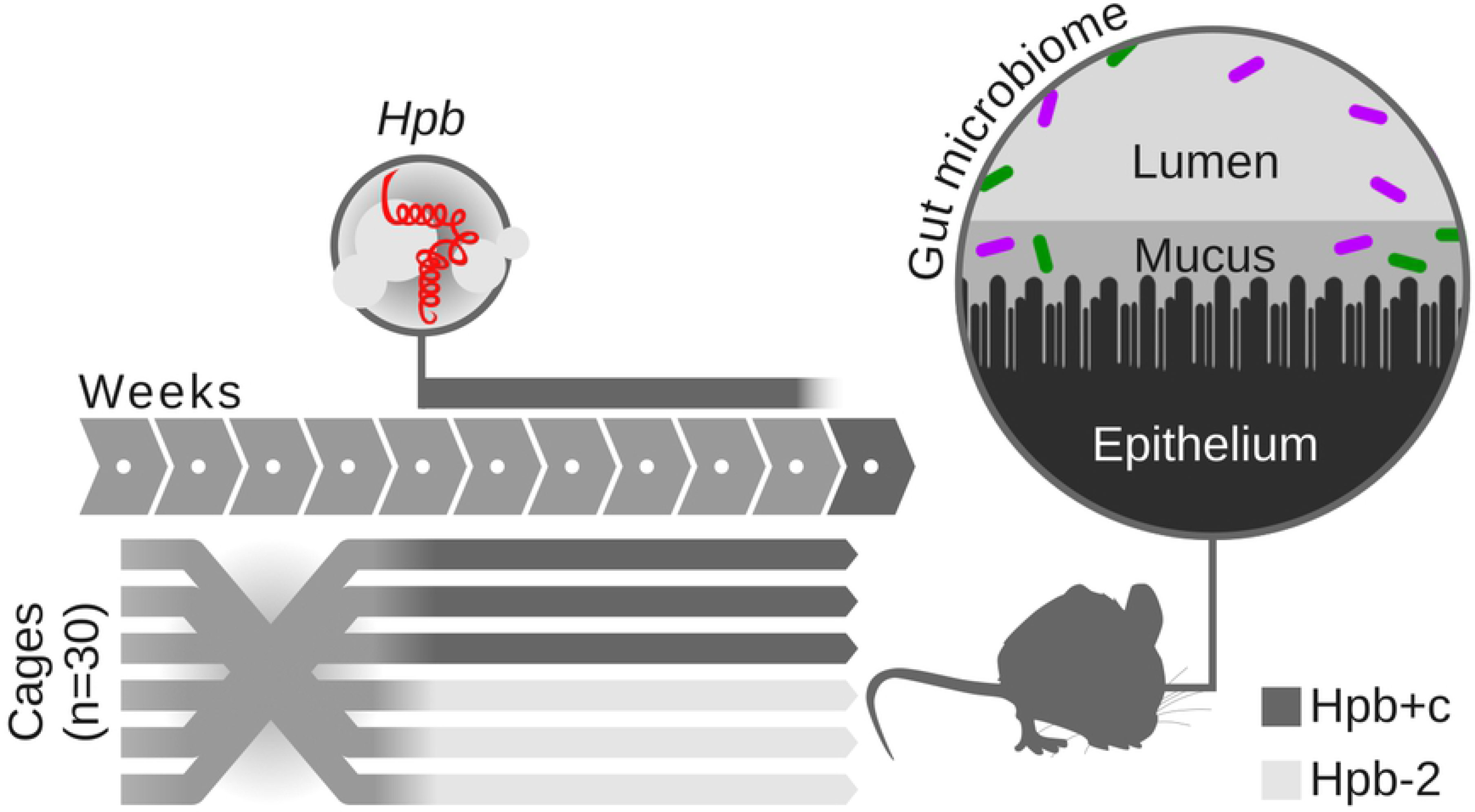
Experimental design of experiment 2 (exp. 2) Analysis of the bacterial communities in the intestinal lumen and intestinal mucus layer 40 days post infection with Hpb larvae.

**Fig 7.**
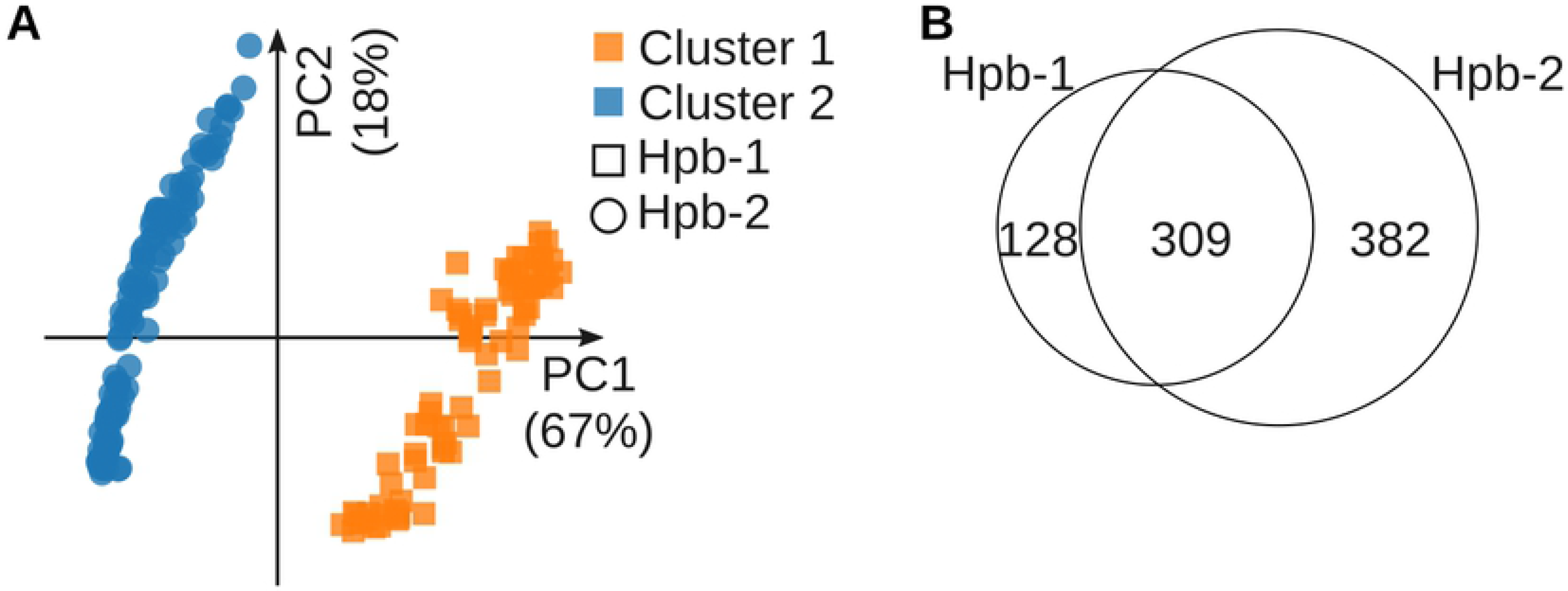
Mice from experiment 1 (exp. 1) and experiment 2 (exp. 2) harbor distinct bacterial communities. (A) Principal coordinates analysis (PCoA) based on the Bray-Curtis dissimilarity for intestinal and fecal bacterial communities (from both mucus layer and lumen samples) in non-infected mice from exp. 1 (Hpb-1) and 2 (Hpb-2). Colors represent clusters of bacterial communities as defined by unbiased similarity network clustering based on Bray-Curtis dissimilarity. (B) Overlaps between non-infected mice from exp. 1 (Hpb-1) and 2 (Hpb-2) for observed bacterial species in intestinal and fecal sample (from the mucus layer and the lumen).

Statistical evidences for an alteration of the microbiome were found in the lumen of the duodenum and in both the mucus layer and the lumen of the ileum, the cecum and the colon (Fig 8A). As seen in exp. 1, a PCoA analysis revealed that intestinal site rather than infection status has the largest impact on bacterial communities structure, with the trend for a clear separation of ileal samples being repeated (Fig 8B). Similarly to infection with larvae batch a in exp. 1, a higher bacterial diversity (measured by the Shannon diversity index) was observed in the lumen of the cecum in infected mice (Fig 8D). In contrast to exp. 1 however, Hpb infection resulted in a lower diversity, both in terms of richness (Fig 8C) and Shannon diversity index (Fig 8D), in the ileum mucus. Differences between Hpb infected and non-infected mice at each sampling site along the intestine notably involved members of the Lachnospiraceae, S24-7, Rikenellaceae, Ruminococcaceae and Peptostreptococcaceae families, two OTUs respectively assigned to orders Clostridiales and Coriobacteriales, an OTU assigned to class Clostridia, members of the Ruminococcus, Turicibacter, Escherichia and Sutterella genera as well as Ruminocuccus gnavus (Fig 9A). As for exp. 1, some bacteria were found significantly (p<0.05) and consistently increased or decreased by the infection across all sampling sites (Fig 9B and Fig 9C). Strikingly, the same bacterium belonging to the Peptostreptococcaceae family that was consistently seen in higher relative abundance in the intestinal mucus layer of infected mice from exp. 1 (Fig 5A and Fig 5C), was also seen consistently higher in both the mucus layer (Fig 9B) and the lumen (Fig 9C) of infected mice in exp. 2, with the exception of the ileal mucus layer where its relative abundance was lower (Fig 9A). In addition, a member of the Clostridiaceae family was found in higher proportions in both intestinal mucus and lumen of infected mice at all sampling sites except the ileal mucus layer, where its relative abundance was lower (Fig 9). As seen in mice infected with larvae from batch b in exp. 1, an OTU assigned to genus Turicibacter was observed to be consistently less abundant along the intestinal lumen in infected mice from exp. 2 (Fig 9). Also, a member of the Lachnospiraceae family together with a member of the genus Sutterella were seen in higher proportions in the lumen at all sampling sites except in the ileum (Fig 9).

**Fig 8.**
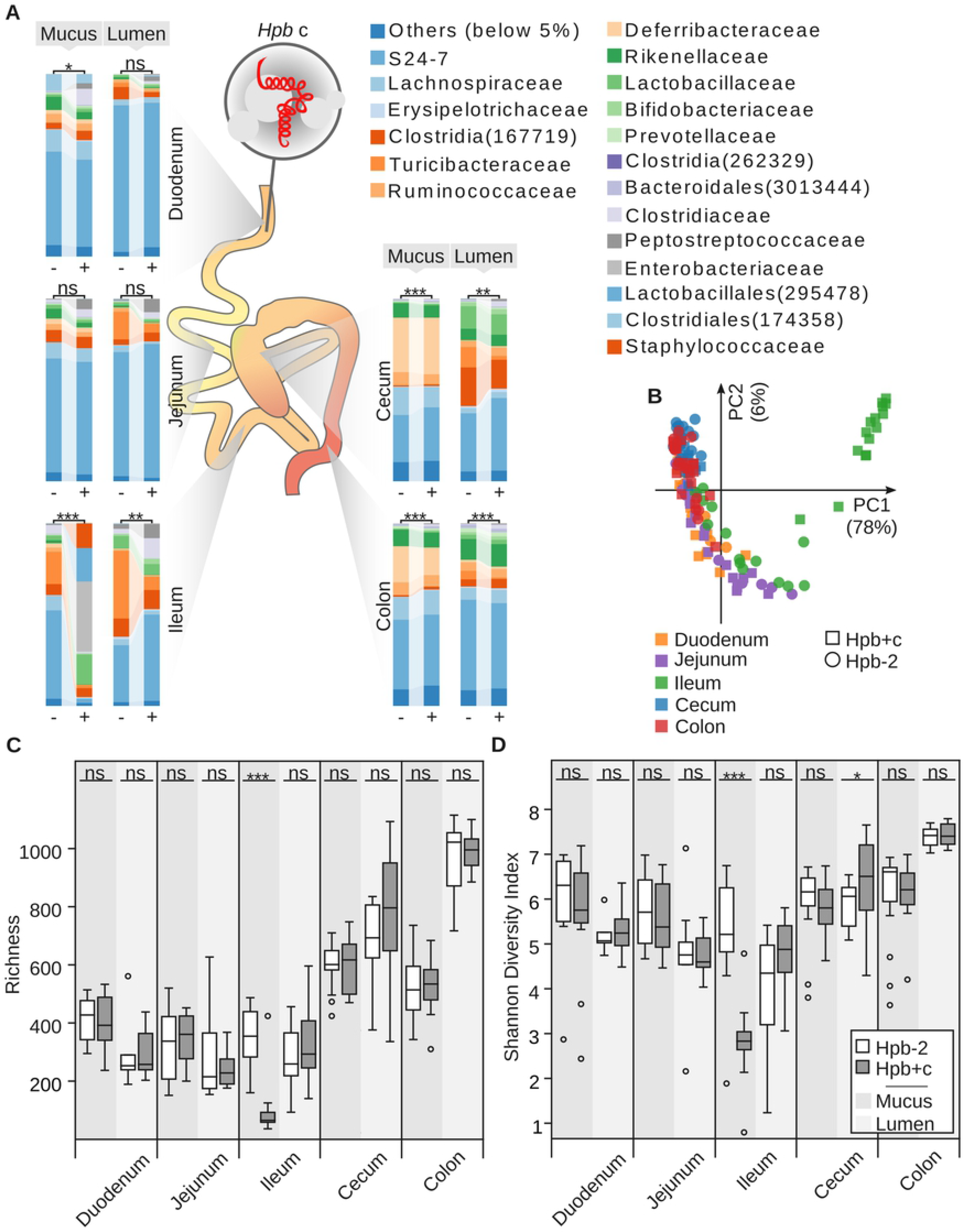
The bacterial community is reproducibly impacted by Hbp infection at multiple sites along the gut, but nature of the impact depends on the initial state of the microbiome. (A) Average proportions of bacterial families in non-infected (-, Hpb-2) and infected (+, Hpb+c) mice from exp. 2. Groups were compared using the Adonis method based on the Jaccard distance (p<0.05: *, p<0.01: **, p<0.001: ***, p>0.05: ns). (B) Principal coordinates analysis (PCoA) based on the Jaccard distance of bacterial communities in the mucus layer along the intestinal tract for non-infected (Hpb-2) and infected (Hpb+c) mice from exp. 2. (C, D) Bacterial communities diversity in the mucus layer and lumen of the intestinal tract and feces for exp. 2 in terms of richness (C) and Shannon diversity index (D). Groups were compared by ANOVA (p<0.05: *, p<0.01: **, p<0.001: ***, p>0.05: ns).

**Fig 9.**
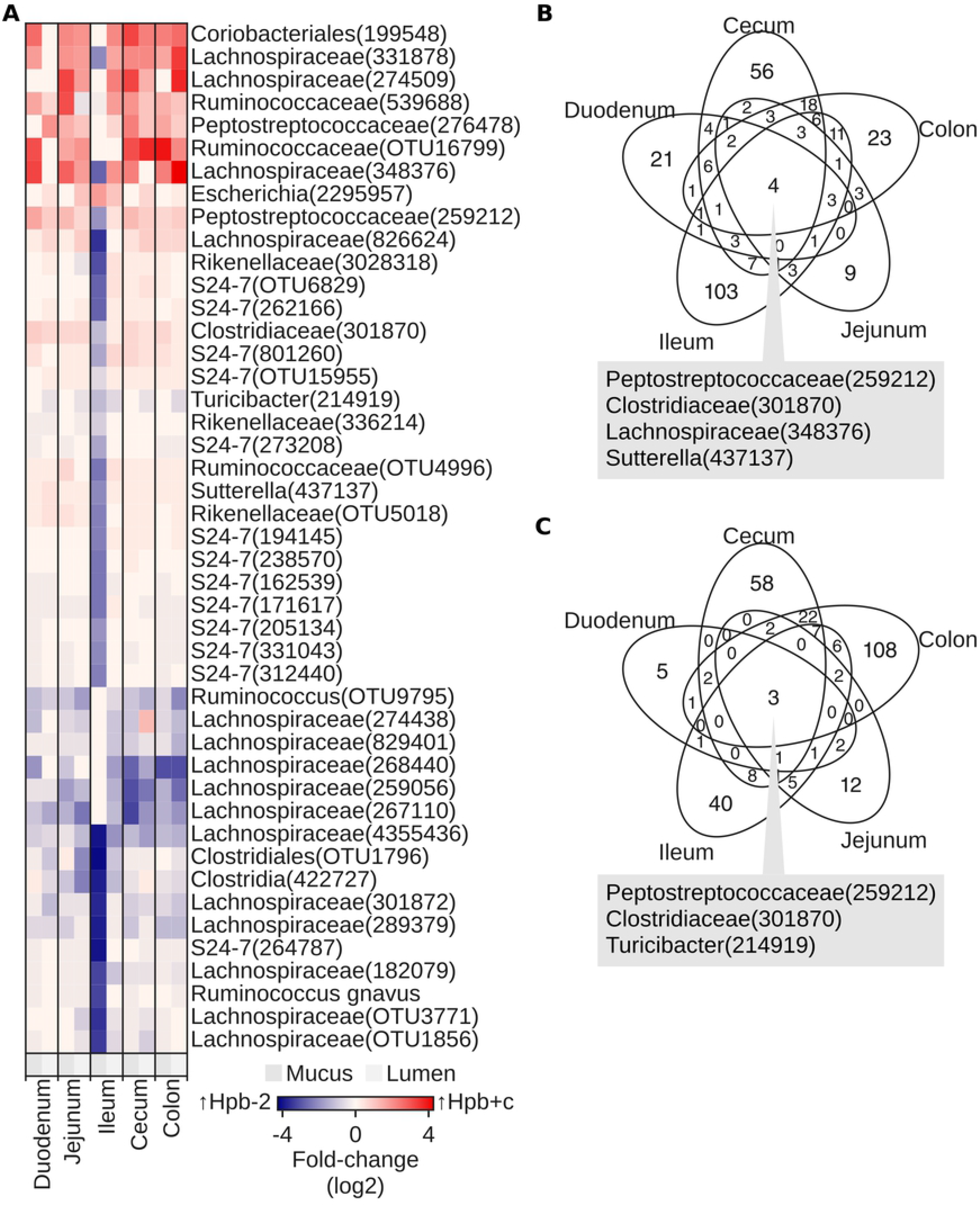
A member of the Peptostreptococcaceae family is consistently associated with Hpb infection across intestinal sites and experimental conditions. (A) As in Fig 4: Differences in proportions of selected bacterial species in the mucus layer and lumen of the intestine and feces between non-infected (Hpb-2) and infected (Hpb+c) mice from exp. 2. (B, C) Overlaps between sampling sites along the intestine for all significant (p<0.05) differences in proportions of observed species between non-infected (Hpb-2) and infected (Hpb+c) mice from exp. 2 in the mucus layer (B) and the lumen (C).

### Helminth-induced alterations to the bacterial microbiome extend well beyond the introduction of new bacterial species by the parasitic larvae

Nematodes are known to carry their own microbiome (24, 25) and Hpb is hatched from laboratory cultures of feces harvested from infected mice and containing parasite eggs (11). We therefore assessed the possibility that bacteria observed in higher proportions within the intestine of infected mice were simply carried there from infecting larvae. To address this question we characterized the microbiome of the same batch of Hpb larvae (batch c) used to perform the infections detailed in exp. 2. Amongst all the bacteria significantly affected by the infection at any sampling site and either in the intestinal mucus layer or in the intestinal lumen, only 26 were detected in Hpb batch c, including Turicibacter and Clostridiaceae (Fig 10A–B). Of note Peptostreptococcaceae 259212 was not present in the Hpb larvae batch c, showing that the effect of Hpb infection on the abundance of this bacterium - which was noted across all experiments performed - was not due to it’s introduction into the host by the parasite.

**Fig 10.**
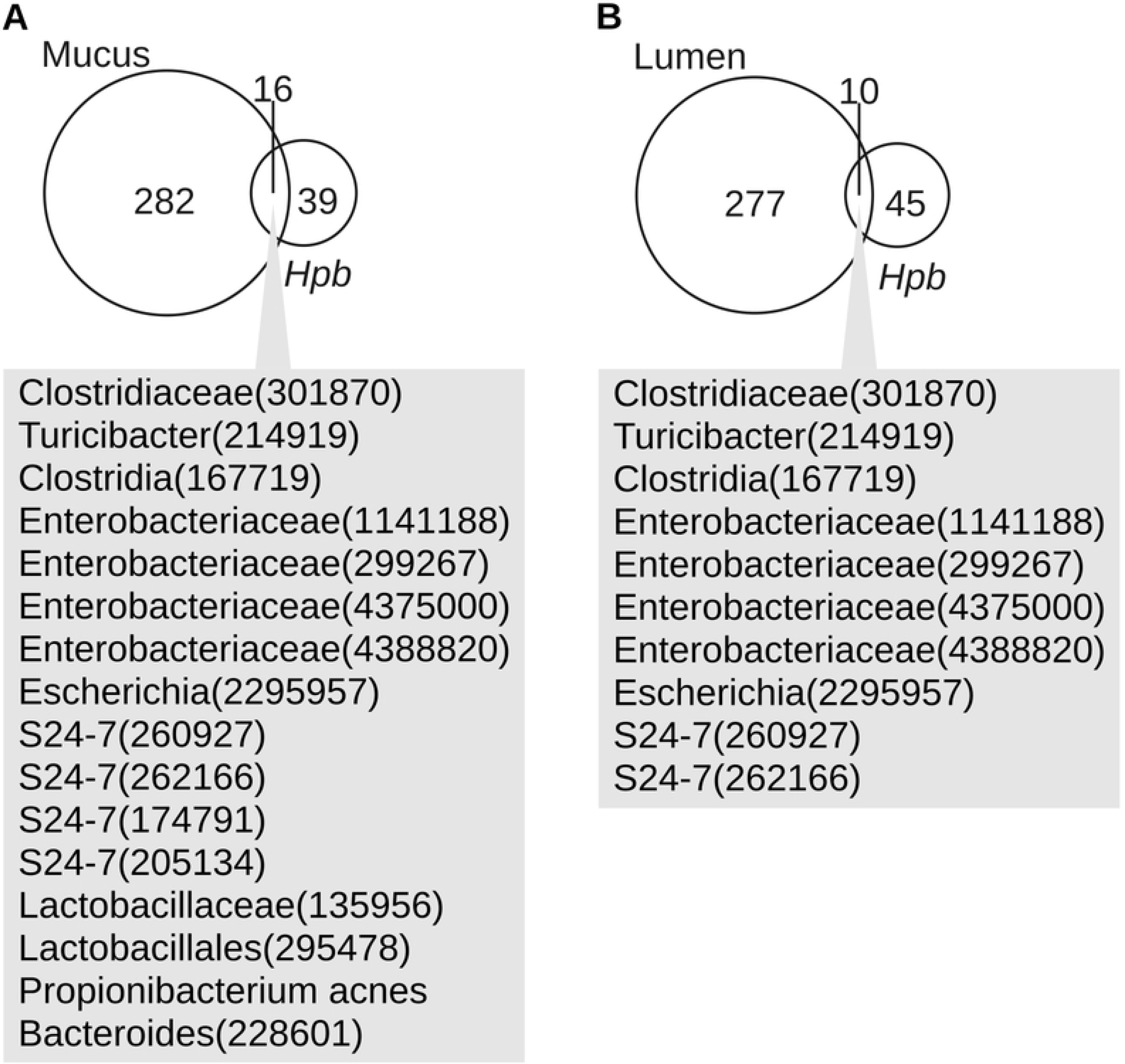
Only a limited number of bacteria found affected by Hpb infection in the gut were observed on Hpb infective larvae before the infection. Venn diagrams highlighting overlaps between bacterial species included in Fig 9B (A) or Fig 9C (B) and bacterial species observed in the batch of infective Hpb larvae used in exp. 2.

## Discussion

The results of this study showed that infection with a small intestinal helminth, Hpb, can consistently and significantly alter bacterial microbial communities throughout the mucous layer and intestinal lumen of the small and large intestine. The use of a large set of mice with a ‘normalized’ microbiome and infected with two distinct batches of helminth larvae provides unequivocal proof that helminth infection can alter the intestinal microbiome. The finding that both the microbiome starting point and the parasite batch employed leads to important differences in the exact nature and site of microbiome alterations is likely to explain, at least in part, why previous studies of helminth-bacterial interactions have reported diverse outcomes. These data highlight the need to carefully control all experiments investigating helminth-microbiome interactions and the difficulty in comparing data generated across different laboratories or even within the same laboratory at different times.

The contribution of parasite batch to variability in helminth-induced microbial alterations likely arises from genetic variations resulting from the sexual life cycle of these complex organisms, however they may also arise from differences in the microbiome associated with the infective larvae. Major differences were also observed in the ‘starting’ microbiome of mice sourced at different times from the same provider, also introducing variability to the data obtained. However, in spite of these variables we found that Hpb infection consistently altered the community structure of bacteria found in the mucous and luminal compartments of the cecum and colon. Alterations also occurred within the small intestine, however the exact compartment affected varied between experimental conditions. In terms of diversity, infection resulted in differences across all experimental conditions but these were smaller and less consistent than alterations to community structure. Of note, variations in bacterial communities along the intestinal tract were poorly reflected in the feces, suggesting a greater stability of the microbial composition in the feces compared to the intestinal tract. This observation emphasizes that a particular attention should be given to sampling site in the study of microbiome. Based on this observation in mice, we further hypothesize that the changes in fecal bacterial composition observed in human after helminths clearance may reflect an even more dramatic change occurring within more distal sites of the intestine.

Although we highlight that different microbiome ‘starting points’ can lead to different responses, metagenomics has shown that a particular niche within the microbiome can be occupied by several species and that communities with different species composition can display more similarity in terms of metabolic functions (26). Therefore, the metagenomic approach used in the present study may fail to capture a more robust impact of Hpb infection at the functional level, a view supported by the previous finding that Hpb consistently leads to higher levels of bacteria-derived SCFAs within the cecum (4). We also consistently noted a higher proportion of a member of the Peptostreptococcaceae family following Hpb infection. This was observed along the entire length of the intestine and across varying experimental conditions. The fact that Hpb larvae did not harbor any Peptostreptococcaceae at the time of the infection rules out the possibility of co-inoculation and indicates that bacteria already present in the host intestine increased in abundance in response to infection. However this observation was only possible because the bacteria in question was already part of the microbial communities prior to infection and may not occur in experiments using mice from other facilities. The Peptostreptococcaceae family (order Clostridiales) includes six described genera which members were isolated from various environments ranging from human and animal microbiome to swine manure and deep-sea hydrothermal vents (27). Of particular interest, this family of bacteria has been described as anaerobic and produces acetate as a product of fermentation. This raises the possibility that the promotion of Peptostreptococcaceae species outgrowth by Hpb may contribute to the previously observed high SCFAs levels during Hpb infection. The 16S rRNA based phylogeny of the Peptostreptococcaceae family has been subject to remodeling and several species of Clostridium have been included into this family in the database of the Ribosomal Database Project (RDP). Based on the alignment of the 16S rRNA gene representative sequence for the Peptostreptococcaceae seen in the present study (OTU 259212) to the NCBI 16S database with BLAST, the phylogeneticly closest identified match was *Romboutsia timonensis* strain DR1 (with only 91% similarity). Altogether, the exact taxonomy of this Peptostreptococcaceae remains unclear in the current state and further characterization (first requiring isolation from the mouse gut) would be necessary in order to understand the possible role of this bacteria within the helminth-associated microbiome and to identify possible mechanisms promoting its outgrowth during Hpb infection.

In conclusion, this work provides conclusive evidence confirming the hypothesis that intestinal helminths can impact on the mammalian intestinal microbiome. Our work also indicates that helminth-induced changes can occur at regions distal to the parasite site of infection and extend well beyond the introduction of new bacterial species carried-over by the infecting larvae. These findings provide impetus for further studies investigating the full impact of helminth-microbial interactions on host health, and determining the molecular mechanisms by which helminths alter microbial communities.

## Materials and Methods

### Ethics statement

All animal experiments were approved by the Service de la consommation et des affaires vétérinaires (1066 Epalinges, Switzerland) with the authorization number 2238.

### Mice

Four weeks old female C57BL/6 wild-type mice were purchased from Charles River Laboratories (France), and housed in specific-pathogen-free (SPF) conditions at the EPFL Faculty of Life Sciences facility for animal housing. Mice were infected with 300 infective L3 Hpb larvae administered by oral gavage in 200ul of saline (Gibco, 10010-015). Control mice received 200ul of saline by the same route.

### Parasites

Hpb L3 infective larvae were generated in the laboratory by Manuel Kulagin and Luc Lebon based on previously described methods (11) Briefly, feces from infected mice were mixed with charcoal and water and incubated at 26°C for one week. The larvae were recovered using a modified Baermann apparatus and washed with saline. Larvae were then incubated for 4 to 6 hours in an antibiotic saline solution containing 5 mg/ml enrofloaxin (injectable Baytril, Bayer), 2 mg/ml amoxicilin and 0.2 mg/ml clavulanic acid (injectable Co-amoxi-Mapha 2200, Mepha Pharma). Larvae were finally washed in saline and stored at 4°C until infection.

### Analysis of intestinal bacterial communities

Bacterial communities were assessed by high-throughput sequencing of the v1-v2 hyper-variable regions of the bacterial 16S rRNA gene as previously described ((28), following the basic protocol “Bacterial 16S rRNA sequencing for bacterial communities present in intestinal contents of mice, or from fecal samples collected from mice or humans”). Samples were processed in nine sequencing runs on an Illumina MiSeq platform using Paired End (PE) v2 2×250 chemistry. The sequences were processed using scripts from the Quantitative Insight Into Microbial Ecology (Qiime) v1.9.0 pipeline (29). Briefly, sequences were trimmed to remove bases showing a Phred quality score lower than 20 using the Seqtk software. Forward and reverse reads were then merged using the join_paired_ends.py script from Qiime (which implements the fastq-join algorithm), setting a minimal overlap of 100bp and allowing a maximal alignment mismatch of 10%. Sample demultiplexing was done using the split_libarries_fastq.py script from Qiime. In addition, reads were truncated at first 3 consecutive bases showing a Phred quality score below 20 and the resulting sequences were discarded if truncated by more than 25% of their length. Reads containing more than 2 N (unknown) nucleotides or were discarded. Reads were finally clustered at 97% similarity and mapped to the GreenGene (30) database as described previously (31). The computation was performed using the clusters of the Vital-it center for high-performance computing of the Swiss Institute of Bioinformatics. Sequences that were not assigned to any taxonomy were discarded. Samples showing less than 5,000 observations were also discarded. The final distribution of samples among experimental groups is describe in Table 1 below.

**Table 1.** Number of samples obtained per intestinal site per experimental group.

|  | Duo.<sup>m</sup> | Duo.<sup>l</sup> | Jej.<sup>m</sup> | Jej.<sup>l</sup> | Ile.<sup>m</sup> | Ile.<sup>l</sup> | Cec.<sup>m</sup> | Cec.<sup>l</sup> | Col.<sup>m</sup> | Col.<sup>l</sup> | Fec. |
| --- | --- | --- | --- | --- | --- | --- | --- | --- | --- | --- | --- |
| Hpb-1 | 4 | 10 | - | 10 | 8 | 8 | 10 | 10 | 10 | 10 | 9 |
| Hpb+a | 6 | 8 | - | 10 | 10 | 6 | 10 | 10 | 9 | 10 | 10 |
| Hpb+b | 5 | 10 | - | 10 | 10 | 10 | 9 | 10 | 10 | 10 | 9 |
| Hpb-2 | 8 | 5 | 10 | 12 | 14 | 15 | 15 | 14 | 15 | 15 | - |
| Hpb+c | 12 | 10 | 14 | 13 | 15 | 15 | 14 | 15 | 15 | 14 | - |

Duo.- duodenum; Jej. - jejunum; Ile. - ileum; Cec. - cecum; Col. - colon; Fec. - feces; <sup>m</sup> – mucus; <sup>l</sup> – lumen.

### Data analysis

Data analysis was performed using Genocrunch (https://genocrunch.epfl.ch). Differences between bacteria communities at each sampling sites in Fig. 2 and Fig. 6.A were assessed at the species level using the Adonis method based on the Jaccard index. Venn diagrams representing significant changes (p<0.05) in individual OTUs across sampling sites were generated in R (32). Rarefaction at the depth of 5,141 observations per sample was applied prior to all analysis with the exception of Fig. 4 and Fig. 7, where rarefaction depth was adapted individually for each sampling site to maximize discovery of bacteria affected by the worm infection (Table 2 below). Differences in individual OTUs proportions and diversity between infected and non-infected mice were assessed by ANOVA. OTUs named in Fig. 4 and Fig. 7.A were selected based on the following criterion: OTUs were first scored based on statistical differences between infected and non-infected groups across sampling sites. Briefly, for each sampling site, a score was assigned (p<0.001 = 3, 0.001<p<0.01 = 2, 0.01<p<0.05 = 1, p>0.05 = 0) and the overall score was calculated as the sum across all sampling site. OTUs showing consistency in fold-change sign at least across five sampling sites together with an overall score higher than five where selected.

**Table 2.** Number of samples obtained per intestinal site per experimental group.

|  | Duo.<sup>m</sup> | Duo.<sup>l</sup> | Jej.<sup>m</sup> | Jej.<sup>l</sup> | Ile.<sup>m</sup> | Ile.<sup>l</sup> | Cec.<sup>m</sup> | Cec.<sup>l</sup> | Col.<sup>m</sup> | Col.<sup>l</sup> | Fec. |
| --- | --- | --- | --- | --- | --- | --- | --- | --- | --- | --- | --- |
| Exp. 1 | 6,213 | 23,528 | - | 17,217 | 26,165 | 42,587 | 53,071 | 9,367 | 20,404 | 49,050 | 17,019 |
| Exp. 2 | 5,318 | 12,306 | 5,058 | 21,777 | 5,168 | 58,390 | 7,204 | 9,955 | 5,873 | 5,442 | - |

Duo.- duodenum; Jej. - jejunum; Ile. - ileum; Cec. - cecum; Col. - colon; Fec. - feces; <sup>m</sup> – mucus; <sup>l</sup> – lumen.

## Acknowledgements

Funding: This work was supported by the European Research Council under the European Union’s Seventh Framework Program (FP/2007-2013)/ERC Grant Agreement [grant number 310948]. The funders had no role in the decision to publish, or the preparation of the manuscript.

## References

1. Rapin A, Harris NL. Helminth-Bacterial Interactions: Cause and Consequence. Trends Immunol. 2018 Jun 22;

2. Maizels RM. Infections and allergy - helminths, hygiene and host immune regulation. Curr Opin Immunol. 2005 Dec;17(6):656–61.

3. Cooper PJ. Interactions between helminth parasites and allergy. Curr Opin Allergy Clin Immunol. 2009 Feb;9(1):29–37.

4. Zaiss MM, Rapin A, Lebon L, Dubey LK, Mosconi I, Sarter K, et al. The Intestinal Microbiota Contributes to the Ability of Helminths to Modulate Allergic Inflammation. Immunity. 2015 Nov 17;43(5):998–1010.

5. Ramanan D, Bowcutt R, Lee SC, Tang MS, Kurtz ZD, Ding Y, et al. Helminth infection promotes colonization resistance via type 2 immunity. Science. 2016 Apr 29;352(6285):608–12.

6. Mamantopoulos M, Ronchi F, Van Hauwermeiren F, Vieira-Silva S, Yilmaz B, Martens L, et al. Nlrp6- and ASC-Dependent Inflammasomes Do Not Shape the Commensal Gut Microbiota Composition. Immunity. 2017 15;47(2):339–348.e4.

7. Bartlett A, Ball PA. Nematospiroides dubius in the mouse as a possible model of endemic human hookworm infection. Ann Trop Med Parasitol. 1972 Mar;66(1):129–34.

8. Gause WC, Urban JF, Stadecker MJ. The immune response to parasitic helminths: insights from murine models. Trends Immunol. 2003 May;24(5):269–77.

9. Pritchard DI, Williams DJ, Behnke JM, Lee TD. The role of IgG1 hypergammaglobulinaemia in immunity to the gastrointestinal nematode Nematospiroides dubius. The immunochemical purification, antigen-specificity and in vivo anti-parasite effect of IgG1 from immune serum. Immunology. 1983 Jun;49(2):353–65.

10. Sukhdeo MV, O’Grady RT, Hsu SC. The site selected by the larvae of Heligmosomoides polygyrus. J Helminthol. 1984 Mar;58(1):19–23.

11. Camberis M, Le Gros G, Urban J. Animal model of Nippostrongylus brasiliensis and Heligmosomoides polygyrus. Curr Protoc Immunol. 2003 Aug;Chapter 19:Unit 19.12.

12. Reynolds LA, Filbey KJ, Maizels RM. Immunity to the model intestinal helminth parasite Heligmosomoides polygyrus. Semin Immunopathol. 2012 Nov;34(6):829–46.

13. Hasnain SZ, Gallagher AL, Grencis RK, Thornton DJ. A new role for mucins in immunity: insights from gastrointestinal nematode infection. Int J Biochem Cell Biol. 2013 Feb;45(2):364–74.

14. Hasnain SZ, Evans CM, Roy M, Gallagher AL, Kindrachuk KN, Barron L, et al. Muc5ac: a critical component mediating the rejection of enteric nematodes. J Exp Med. 2011 May 9;208(5):893–900.

15. Shea-Donohue T, Sullivan C, Finkelman FD, Madden KB, Morris SC, Goldhill J, et al. The role of IL-4 in Heligmosomoides polygyrus-induced alterations in murine intestinal epithelial cell function. J Immunol Baltim Md 1950. 2001 Aug 15;167(4):2234–9.

16. Walk ST, Blum AM, Ewing SA-S, Weinstock JV, Young VB. Alteration of the murine gut microbiota during infection with the parasitic helminth Heligmosomoides polygyrus: Inflamm Bowel Dis. 2010 Nov;16(11):1841–9.

17. Rausch S, Held J, Fischer A, Heimesaat MM, Kühl AA, Bereswill S, et al. Small Intestinal Nematode Infection of Mice Is Associated with Increased Enterobacterial Loads alongside the Intestinal Tract. Allen IC, editor. PLoS ONE. 2013 Sep 10;8(9):e74026.

18. Reynolds LA, Smith KA, Filbey KJ, Harcus Y, Hewitson JP, Redpath SA, et al. Commensal-pathogen interactions in the intestinal tract: Lactobacilli promote infection with, and are promoted by, helminth parasites. Gut Microbes. 2014 Jul;5(4):522–32.

19. Su C, Su L, Li Y, Long SR, Chang J, Zhang W, et al. Helminth-induced alterations of the gut microbiota exacerbate bacterial colitis. Mucosal Immunol. 2018 Jan;11(1):144–57.

20. Rausch S, Midha A, Kuhring M, Affinass N, Radonic A, Kühl AA, et al. Parasitic Nematodes Exert Antimicrobial Activity and Benefit From Microbiota-Driven Support for Host Immune Regulation. Front Immunol. 2018;9:2282.

21. Bouchery T, Volpe B, Shah K, Lebon L, Filbey K, LeGros G, et al. The Study of Host Immune Responses Elicited by the Model Murine Hookworms Nippostrongylus brasiliensis and Heligmosomoides polygyrus. Curr Protoc Mouse Biol. 2017 Dec 20;7(4):236–86.

22. Rausch P, Basic M, Batra A, Bischoff SC, Blaut M, Clavel T, et al. Analysis of factors contributing to variation in the C57BL/6J fecal microbiota across German animal facilities. Int J Med Microbiol IJMM. 2016 Aug;306(5):343–55.

23. Franklin CL, Ericsson AC. Microbiota and reproducibility of rodent models. Lab Anim. 2017 Mar 22;46(4):114–22.

24. Hsu SC, Johansson KR, Donahue MJ. The bacterial flora of the intestine of Ascaris suum and 5-hydroxytryptamine production. J Parasitol. 1986 Aug;72(4):545–9.

25. Zhang F, Berg M, Dierking K, Félix M-A, Shapira M, Samuel BS, et al. Caenorhabditis elegans as a Model for Microbiome Research. Front Microbiol. 2017;8:485.

26. Franzosa EA, Morgan XC, Segata N, Waldron L, Reyes J, Earl AM, et al. Relating the metatranscriptome and metagenome of the human gut. Proc Natl Acad Sci U S A. 2014 Jun 3;111(22):E2329–2338.

27. Slobodkin A. The Family Peptostreptococcaceae. In: Rosenberg E, DeLong EF, Lory S, Stackebrandt E, Thompson F, editors. The Prokaryotes: Firmicutes and Tenericutes [Internet]. Berlin, Heidelberg: Springer Berlin Heidelberg; 2014 [cited 2018 Nov 27]. p. 291–302. Available from: https://doi.org/10.1007/978-3-642-30120-9_217

28. Rapin A, Pattaroni C, Marsland BJ, Harris NL. Microbiota Analysis Using an Illumina MiSeq Platform to Sequence 16S rRNA Genes. Curr Protoc Mouse Biol. 2017 Jun 19;7(2):100–29.

29. Caporaso JG, Kuczynski J, Stombaugh J, Bittinger K, Bushman FD, Costello EK, et al. QIIME allows analysis of high-throughput community sequencing data. Nat Methods. 2010 May;7(5):335–6.

30. DeSantis TZ, Hugenholtz P, Larsen N, Rojas M, Brodie EL, Keller K, et al. Greengenes, a chimera-checked 16S rRNA gene database and workbench compatible with ARB. Appl Environ Microbiol. 2006 Jul;72(7):5069–72.

31. Caporaso JG, Lauber CL, Walters WA, Berg-Lyons D, Huntley J, Fierer N, et al. Ultra-high-throughput microbial community analysis on the Illumina HiSeq and MiSeq platforms. ISME J. 2012 Aug;6(8):1621–4.

32. R Core Team. R: A Language and Environment for Statistical Computing [Internet]. R Foundation for Statistical Computing; 2018. Available from: https://www.R-project.org/

